# Single-cell transcriptome analysis during cardiogenesis reveals basis for organ level developmental anomalies

**DOI:** 10.1101/365734

**Authors:** T. Yvanka de Soysa, Sanjeev S. Ranade, Satoshi Okawa, Srikanth Ravichandran, Yu Huang, Hazel T. Salunga, Amelia Schricker, Antonio Del Sol, Casey A. Gifford, Deepak Srivastava

## Abstract

Organogenesis involves integration of myriad cell types with reciprocal interactions, each progressing through successive stages of lineage specification and differentiation. Establishment of unique gene networks within each cell dictates fate determination, and mutations of transcription factors that drive such networks can result in birth defects. Congenital heart defects are the most common malformations and are caused by disruption of discrete subsets of progenitors^1–3^, however, determining the transcriptional changes in individual cells that lead to organ-level defects in the heart, or other organs, has not been tractable. Here, we employed single-cell RNA sequencing to interrogate early cardiac progenitor cells as they become specified during normal and abnormal cardiogenesis, revealing how dysregulation of specific cellular sub-populations can have catastrophic consequences. A network-based computational method for single-cell RNA-sequencing that predicts lineage specifying transcription factors^4,5^ identified *Hand2* as a specifier of outflow tract cells but not right ventricular cells, despite failure of right ventricular formation in *Hand2*-null mice^6^. Temporal single-cell transcriptome analysis of *Hand2*-null embryos revealed failure of outflow tract myocardium specification, whereas right ventricular myocardium differentiated but failed to migrate into the anterior pole of the developing heart. Dysregulation of retinoic acid signaling, responsible for anterior-posterior patterning^7^, was associated with posteriorization of anterior cardiac progenitors in *Hand2-*null mutant hearts and ectopic atrial gene expression in outflow tract and right ventricle precursors. This work reveals transcriptional determinants in individual cells that specify cardiac progenitor cell fate and differentiation and exposes mechanisms of disrupted cardiac development at single-cell resolution, providing a framework to investigate congenital heart defects.

---

Decades of lineage tracing and loss-of-function studies identified transcriptional regulators required for cardiac differentiation and morphogenesis. The overarching theme from these studies is that gene regulatory network disruption in distinct subsets of cells is devastating for cardiogenesis, despite the relatively normal development of the remaining heart. This observation is consistent with, and begins to explain, clinical manifestations of congenital heart disease (CHD) where discrete regions of the heart are malformed^1–3^. The range of defects is broad, reflecting the numerous cell types involved, and the high susceptibility to perturbation makes CHD the most common birth defect worldwide^1^. Human mutations in essential transcription factors and signaling proteins point to key nodes in regulating distinct heart regions but understanding how gene regulatory network disruption in individual cells contributes to CHD remains unknown. Early efforts to map cardiogenesis at the single-cell level by sequencing hundreds of cells were consistent with recognized heterogeneity of the major cell types^8–10^. However, to gain sufficient resolution to interrogate smaller pools of cellular subtypes required for cardiogenesis, and to investigate gene network disruption in those cell subsets, it may be necessary to study tens of thousands of cells, recognizing that only 1-2 % are specifically disrupted in disease. We addressed this need with droplet-based single-cell RNA-sequencing (scRNAseq), which allowed construction of an unbiased high-resolution molecular framework of early normal and abnormal cardiogenesis.

The heart develops from diverse cell lineages specified from two distinct pools of cardiac progenitor cells (CPCs): the first heart field (FHF) and second heart field (SHF)^1–3^. To identify transcriptional drivers of cardiac cell fate determination and morphogenesis at the single-cell level, we sequenced cells from the cardiogenic region in mouse embryos at three developmental stages: 1) as CPCs begin to differentiate and form a late cardiac crescent at embryonic day (E) 7.75; 2) as the FHF forms a linear heart tube that begins beating and the SHF migrates into the anterior and posterior poles of the tube (E8.25); and 3) as the heart tube loops and incorporates the SHF-derived right ventricle (RV) and atrium with the FHF-derived left ventricle (LV) (E9.25) (**Fig. 1a**). We sequenced the transcriptomes of 38,966 cells, including ectoderm and endoderm progenitors. We computationally excluded non-mesodermal lineages, except neural crest-derived cells that interact with SHF cells, and applied a graph-based clustering approach that partitioned the remaining 21,070 cells into 19 populations (**Fig. 1b, c; Extended Data Fig. 1a, b, Supplementary Table 1**)^11,12^. To assign population identities, we cross-referenced the most highly and uniquely expressed genes in each population with known cardiac subtype markers and *in situ* hybridization data from the literature (**Fig. 1d; Extended Data Fig. 1c, d; Supplementary Table 2**).

**Figure 1:**
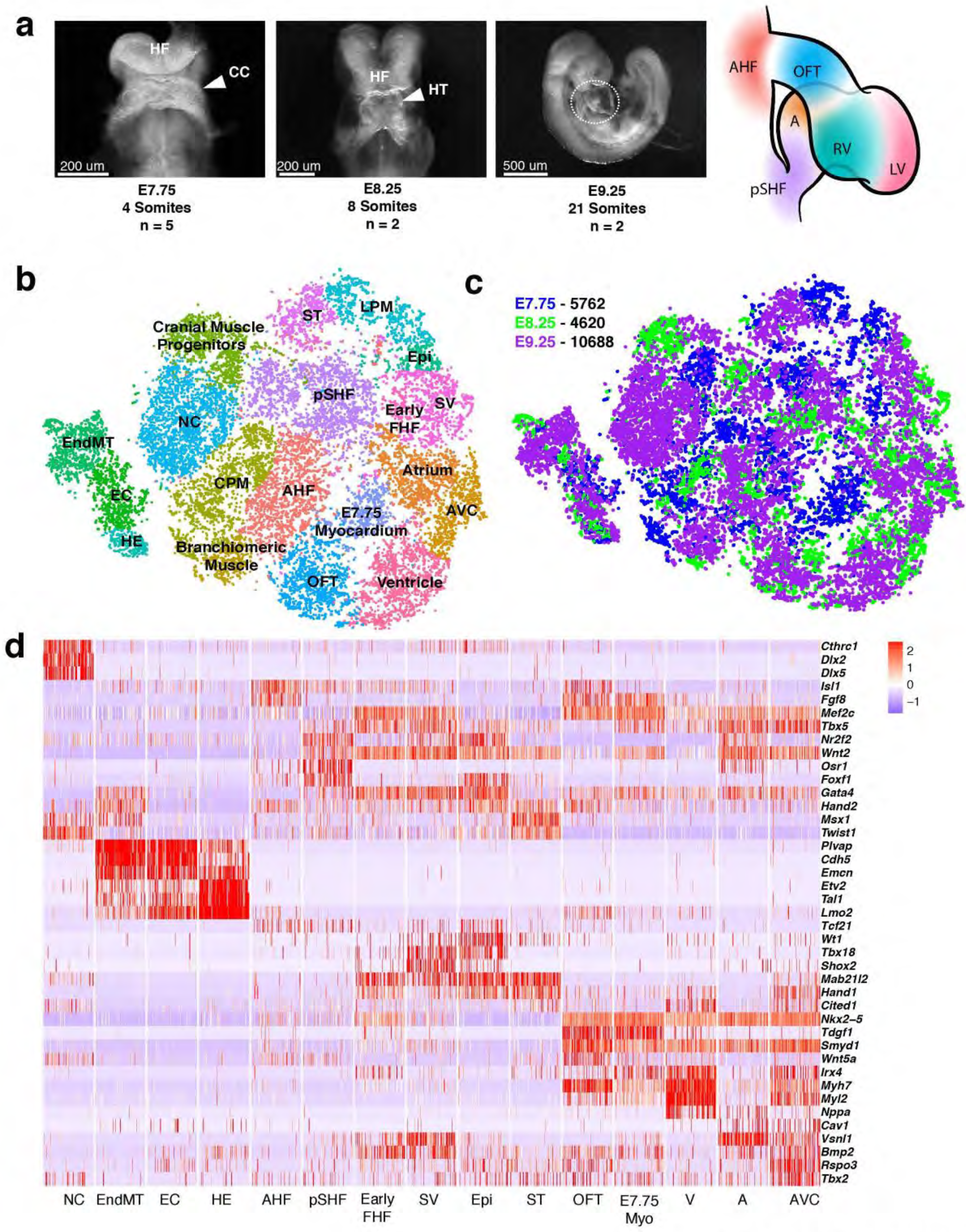
Single-cell RNA-seq reveals heterogeneity of mesoderm- and neural crest-derived lineages. **a**, Representative images of mouse embryos at E7.75, E8.25 and E9.25 used for cell collection. First and second panels indicate frontal views, third panel is right lateral view. Schematic indicates cardiac structures from circled region in third panel. Embryos were somite matched to control for developmental timing heterogeneity. CC, Cardiac Crescent; HF, Head Fold; HT, Heart Tube. **b**, tSNE plot of all mesodermal and neural crest populations captured at E7.75, E8.25 and E9.25 colored by cluster identity. **c**, tSNE plot of all mesodermal and neural crest populations captured at E7.75, E8.25 and E9.25 colored by embryonic stage of collection. Number of cells assayed at each stage is indicated. **d**, Curated expression heatmap of top three highly and uniquely expressed genes of cardiac-relevant populations identified through marker analysis and literature review. Data are shown for 100 cells subsampled from each population. Extra cardiac mesoderm-derived cell types that were captured included the non-cardiac lateral plate mesoderm (LPM), cardiopharyngeal mesoderm (CPM), CPM-derived branchiomeric muscle and cranial muscle. ST, Septum Transversum; Early FHF, Early First Heart Field; pSHF, posterior Second Heart Field; AHF, Anterior Heart Field; Epi, Epicardium; SV, Sinus Venosus; A, Atrium; AVC, Atrioventricular Canal; V, Ventricle; OFT, Outflow Tract; E7.75 myocardium, mixed population of E7.75 myocardial cells; NC, Neural Crest; HE, Hematoendothelial progenitors; EC, Endocardium; EndMT, Endocardium undergoing Endothelial-to-Mesenchymal Transition.

Our clustering analysis revealed that we captured transcriptomes characteristic of the FHF, the anterior and posterior heart field domains of the SHF (AHF and pSHF, respectively), and the cardiomyocyte subtypes specified from these CPC sources (ventricular, outflow tract, atrial, atrioventricular, and sinus venosus). We detected three subpopulations of the endocardial/endothelial lineage: hematoendothelial precursors, specified endothelial/endocardial cells and endocardial cells undergoing an endothelial-to-mesenchymal transition typical of valve development (**Fig. 1d**). We also identified non-cardiomyocyte cell lineages that contribute to cardiac morphogenesis, including epicardial cells and septum transversum cells, from which epicardial cells are derived (**Fig. 1b, d**)^13^. Intriguingly, this analysis revealed a heterogeneous population (labelled as E7.75 myocardium), comprising cells largely captured at E7.75 that expressed the Nodal co-receptor *Tdgf1*^14^ and lower levels of sarcomeric genes such as *Myh7* and *Myl2* compared to outflow tract and ventricular myocardium populations at the later stages (**Fig. 1b, c, d; Extended Data Fig. 1d**). Upon further analysis, this population could be divided into five distinct subpopulations (**Extended Data Fig. 1e, f; Supplementary Table 2**), which we identified as AHF progenitors (A; *Isl1+/Fgf8+/Tdgf1+/Myh7^low^)*, E7.75 RV cells *(B; Irx4+/Myh7+/Hand1-)* and E7.75 LV cells *(C; Irx4+/Myh7+/Hand1+)*, a mix of OFT and RV cells (*D*, Isl1^low^/*Fgf8+/Wnt5a+/Irx4^low^*) and FHF progenitors (*E; Isl1-/Hand1+/Tbx5+)*. This *Isl1-* FHF population was distinct from the *Isl1+* early FHF cells that appear as a unique population in our initial analysis (**Extended Data Fig. 1a, d**) and expressed higher levels of genes such as *Myh6, Tdgf1* and *Nkx2-5*, indicating that it represents a more differentiated FHF progenitor state (**Extended Data Fig. 1g; Supplementary Table 2**). Thus, our data describe previously inaccessible transcriptomes and cell states, representing a rich resource for interrogating the molecular events underlying specification of multiple cardiac subtypes.

Previous studies employing clonal genetic fate mapping and scRNAseq of the earliest *Mesp1+* FHF- and SHF-fated cells demonstrated that these cells are already specified into committed CPCs fated for distinct anatomic regions and lineages between E6.25 and E7.5^15–17^. To determine whether there was additional heterogeneity within the CPC domains, we focused on the early FHF, AHF, and pSHF populations captured at E7.75 and E8.25 (**Extended Data Fig. 1a**), and this analysis further subdivided the CPCs into eight populations (**Fig. 2a, Supplementary Table 2**). In addition to canonical AHF and pSHF cells, we identified a second AHF (AHF2) and two pSHF (pSHF2, pSHF3) populations. AHF2 and pSHF2 were derived from E7.75 and >50% of cells in each population co-expressed the left sided genes *Nodal, Lefty2* and *Pitx2*,^18^ suggesting that these populations contained left-right asymmetrically patterned progenitor cells (**Fig. 2b,c**). Differential gene expression tests between *Pitx2*-positive (normalized UMI > 0.1) and *Pitx2-*negative (normalized UMI < 0.1) cells in AHF2 and pSHF2 identified genes with putative asymmetric expression associated with these left-right asymmetric markers (**Supplementary Table 2**). The pSHF3 population was of unknown specification and was enriched in expression of genes such as *Hoxd1, Hand1, Krt8, Msx1* and *Twist1* and represents a novel progenitor population that will require lineage tracing to determine its contribution (**Fig. 2a, b, c**). Strikingly, we found three distinct *Tnnt2+*/*Isl1*+ double positive populations likely representing the earliest transcriptional states of specified LV CMs, OFT/RV CMs and atrial CMs from the early FHF, AHF, and pSHF, respectively (**Fig. 2a, b**). Pairwise differential gene expression tests identified several genes previously unrecognized to be relatively specific for these cardiac subtype precursors (e.g., *Arl4a, Vegfc* and *Upp1* (atrial), *Nav1, Spry1* and *Dusp14* (OFT/RV), and *Kcng2* (LV) (**Fig. 2c; Supplementary Table 2**).

**Figure 2:**
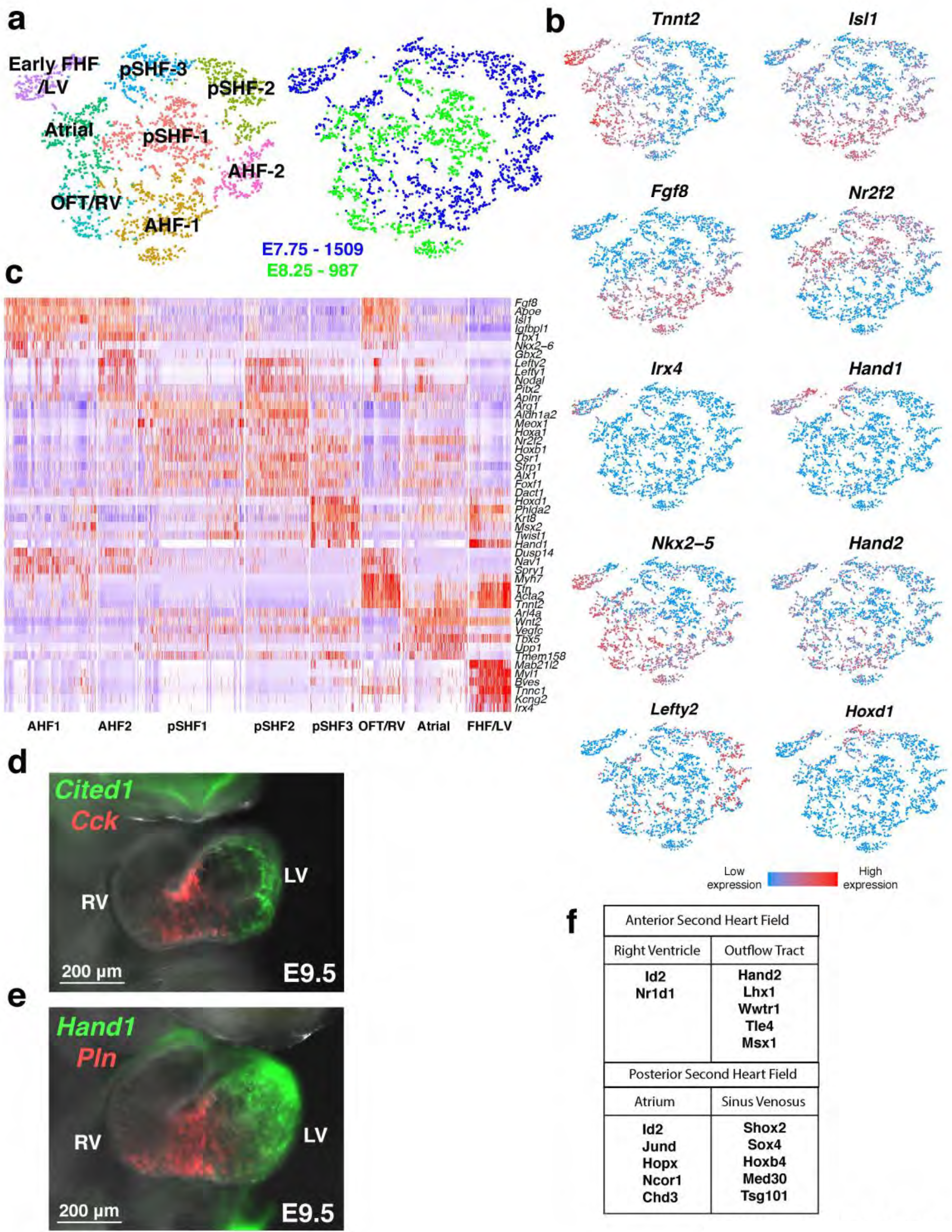
Analysis of cardiac progenitor cell populations reveals early specification dynamics of myocardial subtypes. **a**, tSNE plot of reclustered early FHF, AHF and pSHF populations from Fig. 1b, colored by cluster identity and by embryonic stage of collection. Number of cells from each stage is indicated. **b**, Expression of indicated genes marking subpopulations on tSNE plot. Red or blue indicates high or low expression, respectively. **c**, Curated heatmap of differentially expressed genes between reclustered CPC subpopulations with red or blue indicating high or low gene expression, respectively. All genes represented have an adjusted *p*-value < 1×10^−4^. **d**, Expression of left ventricle marker *Cited1* and newly identified right ventricle gene, *Cck* by whole mount *in situ* hybridization. **e**, Expression of left ventricle marker *Hand1* and *Pln* showing enrichment in right ventricle region by whole mount *in situ* hybridization. **f**, Predicted lineage specifier of OFT and RV cells from the AHF or atrial and sinus venosus cells from the pSHF.

We subdivided the ventricle (V) population (**Fig. 1b**) comprising 1,348 cells to LV and RV, based on the enrichment of LV markers, *Hand1* and *Cited1*, at E8.25 and E9.25 **(Extended Data Fig. 2a, b)**^19^. Several genes were enriched in either chamber, consistent with their unique physiology, although their physiological significance is unclear (**Extended Data Figure 2b, c**; **Supplementary Table 2**). Notably, phospholamban (*Pln*), a critical regulator of calcium handling, and cholecystokinin (*Cck)*, encoding an intestinal hormone, were predominantly expressed in RV cells **(Fig. 2d, e; Extended Data Fig. 2c)**. In situ hybridization confirmed the RV dominance of these genes and also revealed enrichment in the region of the future ventricular septum. These data demonstrate the power of leveraging large numbers of single-cell transcriptomes to reveal unique genes that characterize diversity among highly similar cell subtypes with distinct physiologies.

As CPCs form the cardiac crescent, induction of cardiac gene expression is driven by FGF and BMP signals secreted from the adjacent anterior endoderm^1^. We hypothesized that there may be other endodermal secreted factors expressed by the FGF- and BMP-secreting cells. To test this, we analyzed the *Sox17+/Foxa2+* endoderm cells captured at E7.75 and identified five subpopulations (**Extended Data Fig. 3a; Supplementary Table 2**). Expression of cardiac inducers *Bmp2*, *Bmp4* and *Fgf8* was enriched in different subpopulations, suggesting that cardiac crescent cells receive combinatorial inductive signals from distinct endodermal cell subpopulations (**Extended Data Fig. 3b**). Cluster 3 appears to represent the cardiogenic anterior visceral endoderm, expressing genes such as the Wnt antagonist *Dkk1* and factors such as *Otx2* and *Gsc*, (**Extended Data Fig. 3c**) revealing the undescribed transcriptome of this, and other, specific endodermal populations^20^. For example, we identified *Wnt5a*, a non-canonical *Wnt* ligand, enriched in cluster 1 that co-expressed *Bmp4* (**Extended Data Fig. 3b**). While *Wnt5a* is expressed and required in the SHF^21^, its expression in the endoderm at E7.75 was unappreciated, as broader transcriptomes of cardiogenic endoderm were not formerly accessible. It is intriguing to consider whether the spatial positioning of these endoderm populations relative to the adjacent cardiac mesoderm may determine specification of diverse myocardium subtypes.

The AHF and pSHF give rise to chamber (RV or atria, respectively) and non-chamber (OFT or AVC/sinus venosus, respectively) myocardial lineages. However, the molecular regulators governing differential specification of these lineages are unknown. To identify such lineage specifiers using scRNA-seq data, we applied a Boolean network-based computational method that systematically predicts cell-fate determinants in a system where a parent progenitor cell differentiates to two daughter cells representing different fates^4,5^ (see Methods) (**Fig. 2f**). The appearance of *Id2* as a specifier of both RV and atrium myocardium from their respective progenitor compartments suggests that it may broadly regulate chamber myocardium specification in the SHF. *Shox2* regulates the sinus venosus fate, in part by repressing *Nkx2-5*^22^, and its identification served as a positive control for this analysis. However, it was curious that *Hand2* was predicted as a lineage specifier for the OFT, but not the RV, as global deletion of *Hand2* causes lethality by E10.5 secondary to severe RV hypoplasia^6^. This phenotype is recapitulated upon conditional deletion of *Hand2* in the SHF, underscoring the requirement of *Hand2* in this progenitor compartment^23^. How RV hypoplasia occurs is unknown, in part due to an inability to access individual progenitor cells that may be disrupted early during specification. To resolve the discrepancy of the predicted lineage-specifying function of *Hand2* in the OFT, but not RV, and the morphologic loss-of-function consequence, we investigated the effects of *Hand2* deletion on the fate and behavior of AHF progenitors using single-cell analyses.

We analyzed transcriptomes of nearly 20,000 control and *Hand2-*null cells captured at E7.75 and E8.25 by first processing all control and *Hand2*-null mesoderm populations and focusing on AHF, OFT, RV, LV and atrial precursor populations (**Fig. 3a, b, c; Extended Data Fig. 4a, b, c**). Differential expression analysis revealed that the AHF, OFT/RV (*Tdgf1+/Irx4+*), and RV precursors (*Irx4+* only) in *Hand2*-null embryos were transcriptionally dysregulated (**Supplementary Table 3**) as early as E7.75 (**Fig. 3d, e; Extended Data Fig. 4d**), well before the emergence of a morphologic defect^6^. Notably, the OFT/RV population showed distinct segregation of CTRL and *Hand2*-null progenitors at E8.25 (**Fig. 3c, circled region**) indicating exacerbation of transcriptional dysregulation (**Extended Data Fig. 4d, Supplementary Table 3**). The chromatin remodeling gene *Smyd1* was downregulated in *Hand2*-null AHF cells at E7.75, consistent with the observation that *Smyd1* loss phenotypically mimics that of *Hand2* loss with respect to RV hypoplasia^24^ (**Fig. 3d**). *Smyd1* regulates *Hand2* expression^24^, and our data suggest a possible feedback regulatory mechanism between these two genes. Similarly, *Rgs5*, which we identified as a marker of the SHF, was downregulated in *Hand2*-null cells at E7.75 and E8.25, confirmed by in situ hybridization, indicating failure of normal SHF progenitor identity (**Fig. 3d, h**).

**Figure 3:**
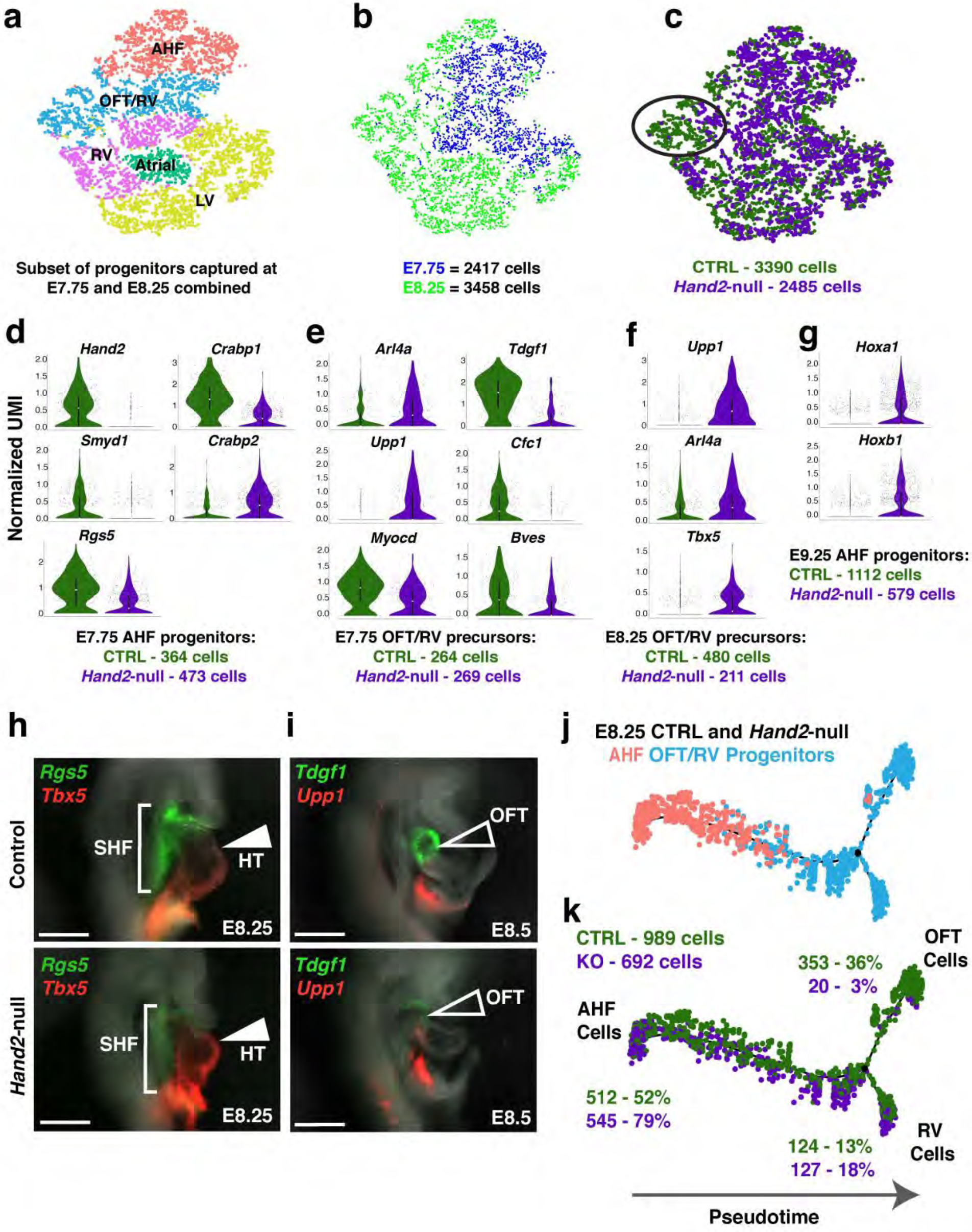
Transcriptional dysregulation in *Hand2-*null embryos reveals AHF specification and differentiation defects with posteriorization of OFT/RV progenitors. **a**, tSNE plot of a subset of E7.75 and E8.25 progenitors relevant to the *Hand2-*null phenotype (OFT, RV, AHF) along with atrial and LV populations colored by cluster. **b**, tSNE plot from Fig. 3a colored by embryonic stage of collection. E7.75 cells are represented in blue and E8.25 cells are represented in light green. **c**, tSNE plot from Fig. 3a colored by genotype. Control (CTRL) data are represented in dark green and *Hand2-*null data are represented in purple. Circled region indicates separation of CTRL and Hand2-null OFT/RV precursors in tSNE space at E8.25. **d**, Violin plots of differentially expressed genes between CTRL and *Hand2-*null AHF progenitors at E7.75. All genes represented have an adjusted p-value < 1×10^−4^. **e**, Violin plots of differentially expressed genes between CTRL and *Hand2-*null OFT/RV progenitors at E7.75. All genes represented have an adjusted *p*-value < 1×10^−4^. **f**, Violin plots of *Upp1*, *Arl4a* and *Tbx5* expression in CTRL and *Hand2-*null OFT/RV progenitors at E8.25 (adjusted *p*-value < 1×10^−4^). **g**, Violin plots of pSHF markers and known retinoic acid target genes *Hoxa1 and Hoxb1* expression in CTRL and *Hand2-*null AHF progenitors at E9.25 (adjusted *p*-value < 1×10^−4^). Summary statistics reported in violin plots: the centre white dot represents median gene expression and the central black rectangle spans the first quartile to the third quartile of the data distribution. The whiskers above or below the box indicate value at 1.5x interquartile range above the third quartile or below the first quartile. **h**, Expression of *Rgs5* and *Tbx5* in CTRL and *Hand2*-null embryos at E8.25 by whole mount *in situ* hybridization. Scale bar indicates 200 μm. **i**, Expression of *Tdgf1* and *Upp1* in CTRL and *Hand2-*null embryos at E8.5 by whole mount *in situ* hybridization. Scale bar indicates 200 μm. **j**, Pseudotime trajectory of AHF and OFT/RV progenitors captured at E8.25 colored by cluster. Black circle indicates branch point where trajectory bifurcates into two lineages **k**, Pseudotime trajectory of AHF and OFT/RV progenitors captured at E8.25 colored by genotype. CTRL, green; *Hand2-*null, purple. Percentages indicate proportion of total cells of one genotype that are from the AHF, OFT or RV lineage. Numbers indicate absolute number of cells of each genotype in each category.

SHF cells undergo a binary decision to adopt an anterior or posterior lineage^17^ and we explored whether this lineage decision is disrupted upon *Hand2* loss. We found that *Crabp1* and *Crabp2* were reciprocally dysregulated in *Hand2-*null AHF cells at E7.75 (**Fig. 3d**). These genes are opposing regulators of retinoic acid (RA) signaling, which is involved in posteriorization of SHF progenitors, resulting in a more atrial fate^7,25^. *Crabp1* is high in the AHF and sequesters RA to facilitate its catabolism, whereas *Crabp2*, which promotes RA nuclear transport and subsequent transcriptional activation^26,27^, is low in AHF cells but high in pSHF cells. In *Hand2-*null AHF cells, *Crabp1* was downregulated, while *Crabp2* was upregulated, which would be expected to result in ectopic RA signaling and posteriorization of the AHF. In agreement with this, early markers of atrial myocardial precursors, *Upp1* and *Arl4a*, were ectopically expressed in *Hand2*-null OFT/RV precursors as early as E7.75 and E8.25, and the pSHF gene *Tbx5*^28^ was ectopically expressed in *Hand2*-null OFT/RV precursors at E8.25 as chambers emerge (**Fig. 3e, f, i**). Strikingly, *Hoxa1* and *Hoxb1*, known RA transcriptional targets^29^ and pSHF markers^17^, were upregulated in *Hand2*-null AHF cells at E9.25 (**Fig. 3g**). These observations suggest that *Hand2* loss results in failure of proper SHF lineage commitment to the AHF, with cells destined to form the OFT and RV developing aberrant features of pSHF cells.

Consistent with AHF cellular identity confusion, genes critical for cardiomyocyte differentiation (e.g., *Tdgf1, Cfc1, Myocd*, *and Bves*)^30,31^, were downregulated in *Hand2-*null OFT/RV precursor cells at E7.75 and E8.25 (**Fig. 3e, i; Supplementary Table 3**). This aberrant gene signature prompted the hypothesis that *Hand2-*null precursors fated to become OFT and RV cells were delayed or disrupted in their differentiation to myocardium. To test this, we ordered control and *Hand2-*null AHF and OFT/RV cells captured at E8.25 in pseudotime, which is a computational measure of the progress a cell makes along a differentiation trajectory^32^. The resulting trajectory began with AHF cells and bifurcated into two distinct lineages (**Fig. 3j, Extended Data Fig. 5a**). We used genes with significant lineage-dependent expression^32^ to classify each lineage as OFT or RV (**Extended Data Fig. 5b**). Strikingly, *Hand2-*null cells were severely depleted in the OFT lineage (**Fig. 3k**). In contrast, *Hand2-*null cells populated the RV lineage in numbers comparable to control cells. These data suggest differential perturbations in OFT- and RV-fated cells upon *Hand2* loss; whereas OFT-fated cells had disrupted specification, RV cells were appropriately specified, consistent with the lineage-specifier analysis. However, this is discrepant with histological evidence that *Hand2*-null embryos have no discernible RV chamber at E8.25 or E9.25^6^. To resolve this and determine the nature of the still-specified *Hand2-*null RV cells, we analyzed control and *Hand2-*null progenitor transcriptomes captured at E9.25 (**Fig. 4a, b**). Expression overlap between the ventricle-specific gene *Irx4* and the newly identified RV-enriched *Cck* gene, as well as exclusion of LV genes *Hand1* and *Cited1*, indicated the presence of two distinct RV populations (**Fig. 4c; Extended Data Fig. 6a**). Notably, each population exclusively comprised control or *Hand2-*null cells, suggesting that RV cells are indeed present in *Hand2-*null hearts at this stage despite the absence of the RV chamber (**Fig. 4b**).

**Figure 4:**
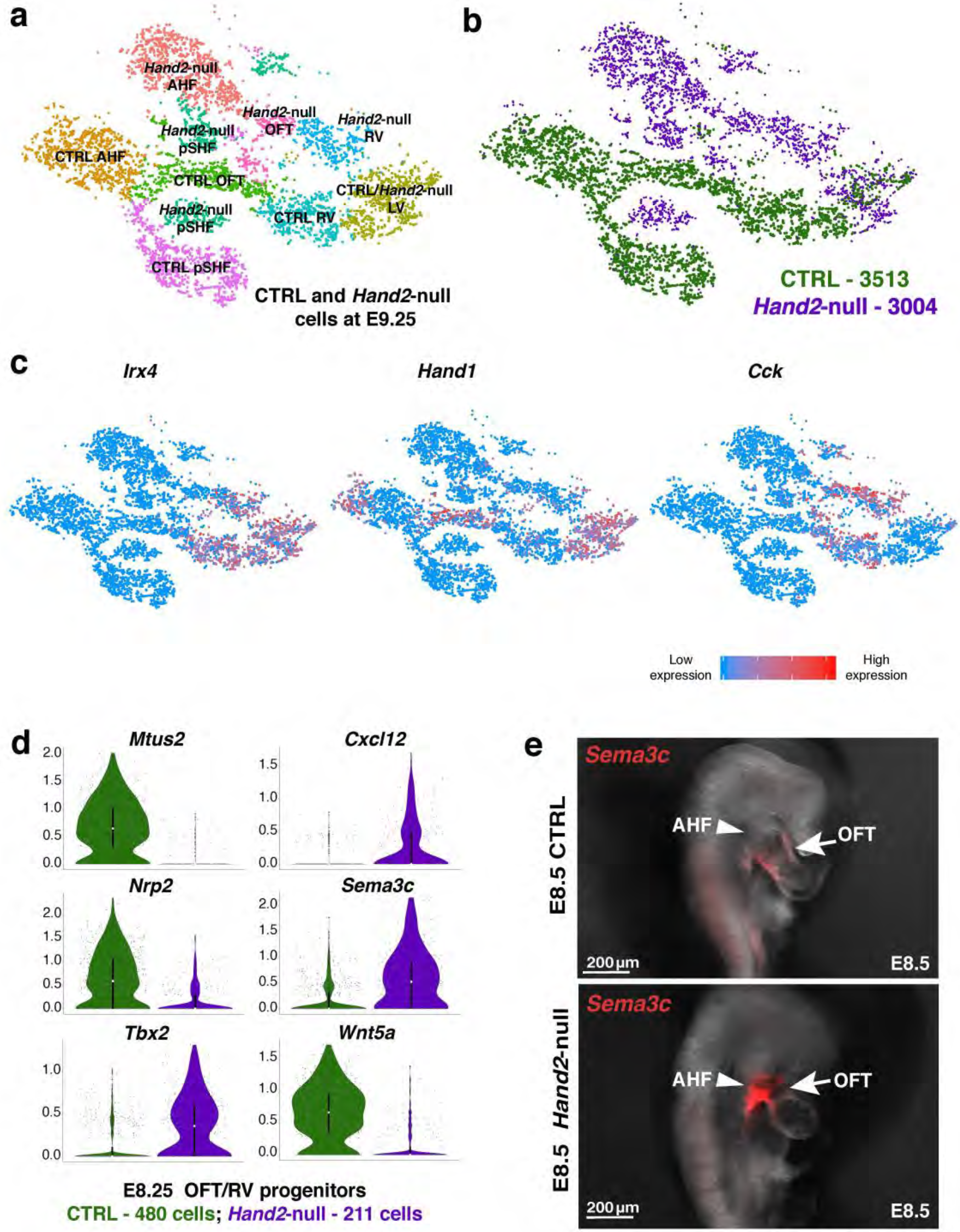
RV cells are appropriately specified but fail to migrate into the developing heart. **a**, tSNE plot of CTRL and *Hand2-*null SHF, Ventricle and OFT populations at E9.25 colored by cluster. **b**, tSNE plot of CTRL and *Hand2-*null SHF, Ventricle and OFT populations at E9.25 colored by genotype. **c**, Expression overlap of *Irx4, Hand1* and *Cck* indicating presence of LV and RV cell populations. Red or blue indicates high or low levels of expression, respectively. **d**, Violin plots showing dysregulation of migration related genes in CTRL and *Hand2-*null OFT/RV progenitors at E8.25. All genes represented have an adjusted *p*-value < 1×10^−4^. Summary statistics reported in violin plots: the centre white dot represents median gene expression and the central black rectangle spans the first quartile to the third quartile of the data distribution. The whiskers above or below the box indicate value at 1.5x interquartile range above the third quartile or below the first quartile. **e**, *In situ* hybridization of *Sema3c* mRNA expression in whole-mount CTRL or *Hand2-*null embryos at E8.5.

To explore the underlying mechanism for defective RV chamber formation, we investigated if RV-specified cells were mis-localized during development. *In-situ* hybridization revealed *Irx4+/Cck*+ cells marking the RV behind the LV in the area of the OFT, suggesting a migratory defect (**Extended Data Fig. 6b**). Consistent with this, *Mtus2, Nrp1* and *Nrp2*, which are involved in cellular migration and guidance cues^33,34^, were severely downregulated in *Hand2-*null E8.25 OFT/RV cells (**Fig. 4d**). In contrast, the chemotactic secreted ligands, *Cxcl12*^35^ and *Sema3c*, were prematurely upregulated in those cells. *Sema3c* is typically expressed in OFT myocardium to attract *PlexinA2*-positive neural crest cells^34^. Strikingly, in *Hand2-*null embryos, *Sema3c*-positive cells accumulated in the pharyngeal mesoderm behind the heart instead of populating the cardiac outflow tract (**Fig. 4e**), in agreement with arrested migration of AHF-derived cells. Moreover, *Tbx2* and *Wnt5a* were ectopically activated or downregulated, respectively, in *Hand2*-null OFT/RV progenitor cells (**Fig. 4d**). Experimental misexpression of *Tbx2* in the AHF and *Wnt5a* deletion have previously been shown to inhibit deployment of AHF cells into the heart tube^36,37^, suggesting their dysregulated expression in *Hand2-*null OFT/RV cells may contribute to the migratory defect.

At E9.25, *Hand1* and *Hand2* are co-expressed in LV progenitors (**Extended Data Fig. 6c**) and *Hand1* is redundant with *Hand2*^38^, consistent with the intact LV in *Hand2*-null embryos. *Hand1* is also co-expressed in OFT progenitors, but curiously fails to compensate for *Hand2* loss in this cell type. The quantitative scRNAseq approach revealed that *Hand1* was selectively downregulated in *Hand2*-null OFT, but not LV, progenitors (**Extended Data Fig. 6c**), suggesting distinct spatial regulatory logic and dependence upon *Hand2*. Thus, *Hand2*-null OFT progenitors lack *Hand1* and *Hand2*, which may explain the failure of OFT specification.

In this study, we demonstrate the power of scRNAseq to reveal regulatory defects in minor subsets of cells, leading to mechanistic understanding of morphologic developmental defects. The combination of scRNAseq and computational network analyses led to predictions that were experimentally validated regarding the essential function of *Hand2* in specifying OFT cells during cardiogenesis. The ability to acquire quantitative and spatial resolution of transcriptomes provided evidence for posteriorization of the AHF progenitors and gene dysregulation resulting in specification and migration defects. Because of the relatively few cells in the heart that were affected in *Hand2* mutants, these features remained undiscovered and likely were revealed by our ability to interrogate tens of thousands of cells in the organ temporally. These features of single-cell transcriptomics offer a powerful strategy to more effectively map the mechanisms underlying developmental defects reported in animal models and thereby reveal causes of related congenital anomalies. This is a prerequisite to preventive approaches for birth defects and potential post-natal intervention for ongoing sequelae of human malformations.

## Materials and Methods

### Mice

All protocols concerning animal use were approved by the IACUC at the University of California San Francisco and conducted in strict accordance with the NIH *Guide for the Care and Use of Laboratory Animals*. Transcriptomes were captured from wildtype (WT), Hand2^+/−^(HET), and *Hand2-*null embryos from intercrossed C57BL/6 mice heterozygous for the *Hand2*-null allele^6^. Since HET embryos were morphologically indistinguishable from WT embryos^6^ and did not show transcriptional dysregulation by scRNAseq, transcriptomes were combined as controls in the *Hand2*-null analyses.

Timed matings were set up where E0.5 was considered as the day of plug detection. Pregnant females were identified by echocardiography performed at E6.5 and sacrificed to harvest embryos at E7.75, E8.25 and E9.25. Transcriptomes from at least 2 embryos were collected per embryonic stage, per genotype. Embryos were developmentally matched at each time point by somite count (4, 8, and 21 somites from E7.75, E8.25, and E9.25, respectively).

### Embryo dissection and single-cell library generation

Embryos were dissected in cold PBS (Life Technologies, CAT# 14190250), de-yolked and placed in PBS/1% FBS (ThermoFisher Scientific, CAT# 10439016) solution on ice until dissociation (approximately 3 hours). Yolk sac DNA was extracted (QuickExtract DNA Extraction Solution, Epicentre, CAT# QE09050) and used for genotyping to distinguish *Hand2* WT, HET and *Hand2-*null embryos before further microdissection of cardiac regions at each stage. Dissected cardiac tissue was incubated in 200 μl TrypLE (ThermoFisher Scientific, CAT# 12563029) for 5 min, triturated with a 200 μl pipette tip, and incubated for an additional 5 min. The TrypLE solution was quenched with 600 μl PBS/1% FBS. Cells were filtered through a 70 μm cell strainer (BD Falcon, CAT# 08-771-2), centrifuged at 150 rcf for 3 min, and resuspended in 35 μl PBS/1% FBS. Single-cell droplet libraries from this suspension were generated in the Chromium controller according to the manufacturer’s instructions in the Chromium Single Cell 3′ Reagent Kit v2 User Guide. Additional components used for library preparation include the Chromium Single Cell 3′ Library and Gel Bead Kit v2 (PN-120237) and the Chromium Single Cell 3′ Chip kit v2 (PN-120236).

### Single-cell transcriptome library preparation and sequencing

Libraries were prepared according to the manufacturer’s instructions using the Chromium Single Cell 3′ Library & Gel Bead Kit v2 (PN-120237) and Chromium i7 Multiplex Kit (PN-120262). Final libraries were sequenced on the NextSeq 500 and Hiseq 4000. Somite-matched WT, HET and *Hand2-*null replicate libraries from each litter were always pooled and sequenced in the same lane. Sequencing parameters were selected according to the Chromium Single Cell v2 specifications. All libraries were sequenced to a mean read depth of at least 50,000 total aligned reads per cell.

### Processing of raw sequencing reads

Raw sequencing reads were processed using the Cell Ranger v1.3.1 pipeline from 10X Genomics. Briefly, reads were demultiplexed, aligned to the mouse mm10 genome and UMI counts were quantified per gene per cell to generate a gene-barcode matrix. Data from multiple samples (WT only analysis, WT/HET/ *Hand2-*null analysis) were aggregated and normalized to the same sequencing depth, resulting in a combined gene-barcode matrix of all samples.

### Cell filtering and cell-type clustering analysis

We sequenced the transcriptomes of 25,512 cells captured from WT, 13,402 cells from *Hand2^+/−^*, and 33,897 cells captured from *Hand2^−/−^* embryos. Further filtering and clustering analyses of these cells were performed with the Seurat v2.2 R package, as described in the tutorials (http://satijalab.org/seurat/)^11^. Cells were normalized for genes expressed per cell and total expression, and cell cycle stage, then multiplied by a scale factor of 10,000 and log-transformed. We used regression to eliminate technical variability due to the number of genes detected, day of library preparation, and stage of the cell cycle. Significant PCs were used for downstream graph-based, semi-unsupervised clustering into distinct populations and used t-distributed Stochastic Neighbor Embedding (tSNE) dimensionality reduction was used to project these populations in a 2D plot. To identify marker genes, the resulting clusters were compared pairwise for differential gene expression using the Likelihood-ratio test for single-cell gene expression (*FindAllMarkers* function). We excluded endoderm- and ectoderm-derived clusters, retaining the cells of mesodermal or neural crest identity for further clustering and analysis. For the WT analysis in Figure 1, data were processed and reclustered using the single-cell alignment procedure^12^. We elected to do this because initial clustering and tSNE visualization showed that populations derived from E9.25 were consistently separated from earlier time points, likely due to technical variation arising from differences in timing of embryo dissection and processing. For all other analyses, combinations of clusters obtained with the *SubsetData* function were processed similarly as described above with regression of technical variables, identification of highly variable genes, principal component analysis, graph-based clustering, tSNE projection and marker analysis. An adjusted *p*-value (Bonferroni Correction) cut-off < 1×10^−4^ was used to identify differentially expressed genes between WT and *Hand2-*null cells (*FindMarkers function*). R scripts are available upon request.

### Prediction of cell fate determinants

Cell fate determinants for OFT and RV from the AHF, and for SV and A from the pSHF were predicted using a modified version of the method that we previously developed^4,5^. One hundred cells were randomly selected from a parental subpopulation (i.e., AHF or pSHF) and from a corresponding daughter subpopulation (i.e., OFT or RV for AHF, SV or A for pSHF) and the normalized ratio difference (NRD) was computed for all combinations of these 100 cells, yielding 10,000 parent-daughter cell combinations. The NRD was calculated for all pairs of differentially expressed TFs between OFT and RV for the AHF differentiation event, and between SV and A for the pSHF differentiation event and averaged over the 10,000 cell combinations. TF pairs whose mean NRD was more than 0.05 in one lineage direction but less than 0.01 in the other lineage direction were selected. Finally, the TF pairs that resided in the strongly connected component of the GRN were kept as the final candidate cell fate determinants.

### Cell trajectory analysis

AHF and OFT/RV progenitors from WT and *Hand2-*null cells captured at E8.25 were ordered in pseudotime with the Monocle 2 package, as described in the tutorials (http://cole-trapnelllab.github.io/monocle-release/)^32^. The differentially expressed genes between these two clusters, as determined in Seurat, were used as input for temporal ordering of these cells along the differentiation trajectory. We used BEAM, which identifies genes with significant branch-dependent expression (adjusted *p*-value < 1×10^−15^), to determine the identities of the two lineages diverging from the trajectory branch-point as OFT and RV.

### *In Situ* hybridization experiments

Each *in situ* hybridization experiment was replicated twice. For whole-mount experiments: deyolked whole embryos were fixed in a 4% formaldehyde solution (ThermoFisher Scientific, CAT# 28906) overnight at 4°C followed by 2X PBST washes and 5-minute incubations in a dehydration series of 25%, 50%, 75% and 100% methanol (Fisher Scientific, CAT# A454-1). At this point embryos were stored in 100% methanol at −20°C until the *in situ* protocol was initiated. Yolk sac DNA was used for genotyping. The whole-mount *in situ* assay was adapted from the protocol formulated for whole-mount zebrafish embryos^39^ using the RNAscope Multiplex Fluorescent Reagent Kit v2 (Advanced Cell Diagnostics, CAT# 323100), with minor modifications (the air-drying step was excluded in our protocol, Protease Plus was used for embryo permeabilization, and the 0.2X SSCT wash step between reagent incubations was reduced to 3X 8 mins). Whole-mount embryos were imaged on the Leica M165 fluorescent dissecting scope.

For *in situ* hybridization experiments performed on embryo sections: embryos were washed 3X in PBS after overnight fixation in 4% formaldehyde and stored in 70% ethanol (VWR, CAT# 89125-186) indefinitely until embedding. Embryos were embedded in Histogel (Thermo Scientific, CAT# R904012) and paraffin processed using standard protocols and embedded for transverse sectioning. Tissue slices were serially sectioned at 5 μm intervals, mounted on slides and stored at room temperature until initiation of the RNAscope protocol for paraffin embedded sections (User manual catalog number 322452-USM).

Catalog numbers for RNAscope probes used in this study: *Cck*, 402271-C3; *Cited1*, 432471; *Hand1*, 429651-C2; *Irx4*, 504831; *Pln*, 506241; *Rgs5*, 430181; *Sema3c*, 441441-C3; *Tdgf1*, 506411; *Tbx5*, 519581-C2 and *Upp1*, 504841-C2.

## Data availability

All source data, including sequencing reads and single-cell expression matrices, will be available from GEO upon publication.

## Supplementary Information

**Supplementary Table 1** – This table summarizes features of analyzed cells at each embryonic stage for each biological replicate of wild-type, *Hand2* heterozygous and *Hand2*-null embryos.

**Supplementary Table 2** – This table lists differentially expressed genes used to identify and distinguish populations discussed in Figure 1, Figure 2, and Extended Data 1, 2, and 3.

**Supplementary Table 3** – This table lists differentially expressed genes between control and *Hand2*-null populations.

## Acknowledgements

The authors thank Dr. Benoit Bruneau and members of the Srivastava lab for helpful discussion and feedback and Dr. Cole Trapnell for guidance on scRNAseq analysis. The authors acknowledge the Gladstone Histology and Light Microscopy Core, the Gladstone Genomics Core and the Gladstone Bioinformatics Core for their technical expertise and the Gladstone Animal Facility for support with mouse colony maintenance. We thank T. Marsh, B. Taylor, T. Roberts and G. Maki for their assistance with imaging, editing and graphics. Y.D.S was supported by the UCSF Chancellor’s Fellowship, Genentech Foundation Fellowship, Discovery Fellows Program and Phi Beta Kappa Graduate Scholarship. C.A.G. is a HHMI fellow of the Damon Runyon Cancer Research Foundation (DRG-2206-14). D.S. is supported by the National Heart Lung and Blood Institute (R01 HL057181, U01 HL098179), the California Institute for Regenerative Medicine (DISC2-09098), the Roddenberry Foundation, the L.K. Whittier Foundation, and the Younger Family Fund. S.O. is supported by an FNR CORE grant (C15/BM/10397420) and S.R by University of Luxembourg IRP Grant (R-AGR-3227-11). This work was also supported by NIH/NCRR grant C06 RR018928 to the Gladstone Institutes.

## Author Contributions

Y.D.S., C.A.G. and D.S. conceived the study, interpreted the data and wrote the manuscript. Y.D.S. prepared chromium libraries, performed *in situ* hybridization experiments and imaging, and analyzed data with Seurat and Monocle. Y.D.S. and S.S.R. dissected and processed embryos for single-cell library preparation and *in situ* hybridization experiments. Y.D.S. and A.S. performed genotyping of mice. S.O., S.R. and A.D.S. conducted computational modeling for cell fate determinant predictions. Y.H. and H.S. identified pregnant female mice by echocardiography.

## Author Information

The authors declare no competing financial or non-financial interests as defined by Nature Research. Correspondence and requests for materials should be addressed to D.S.

---

**Extended Data Figure 1:**
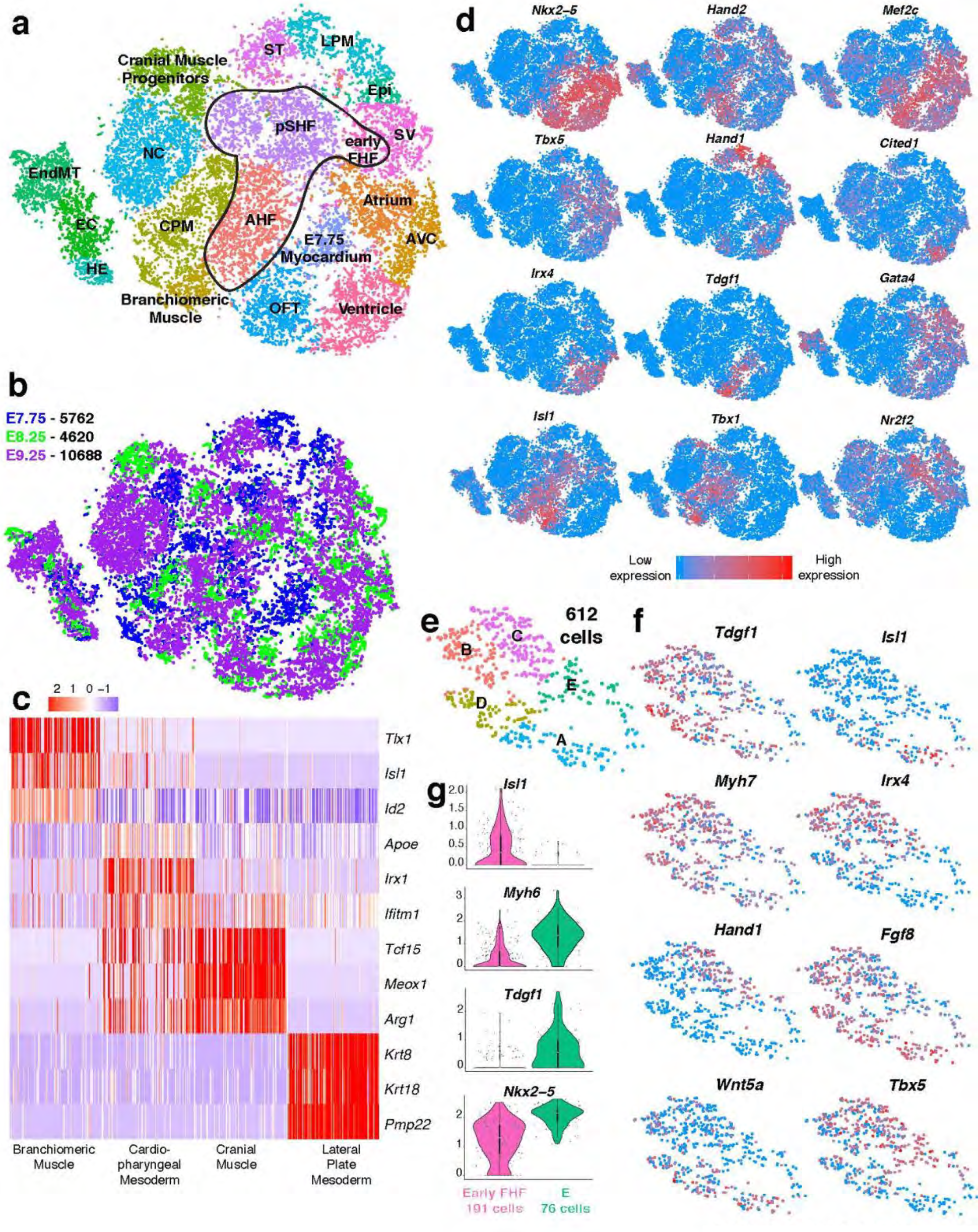
Expression patterns of known marker genes in all captured populations. **a**, tSNE plot of captured mesoderm and neural crest lineages from Fig. 1b colored by cluster. Circled region indicates populations that were reclustered and depicted in Fig. 2. **b**, tSNE plot of captured mesoderm and neural crest lineages from Fig. 1b colored by embryonic stage of collection. **c**, Expression heatmap of top three highly expressed genes of non-cardiac populations identified through marker analysis and literature review. Data are shown for 100 cells subsampled from each population. **d**, Expression pattern of cardiac transcription factors and signaling proteins that define cardiac cell lineages. LPM, Lateral Plate Mesoderm; CPM, cardiopharyngeal mesoderm; BMsc, Branchiomeric Muscle; CMsc, Cranial Muscle. **e**, tSNE plot of reprocessed cells from E7.75 myocardium cluster in Extended Data Fig. 1a. **f**, Expression of key marker genes of subpopulations depicted in tSNE in Extended Data Fig. 1e. **g**, Violin plots showing expression of *Isl1, Myh6, Tdgf1* and *Nkx2-5* in early FHF and FHF cells in population E from Extended Data Fig. 1e. Summary statistics reported in violin plots: the centre white dot represents median gene expression and the central black rectangle spans the first quartile to the third quartile of the data distribution. The whiskers above or below the box indicate value at 1.5x interquartile range above the third quartile or below the first quartile.

**Extended Data Figure 2:**
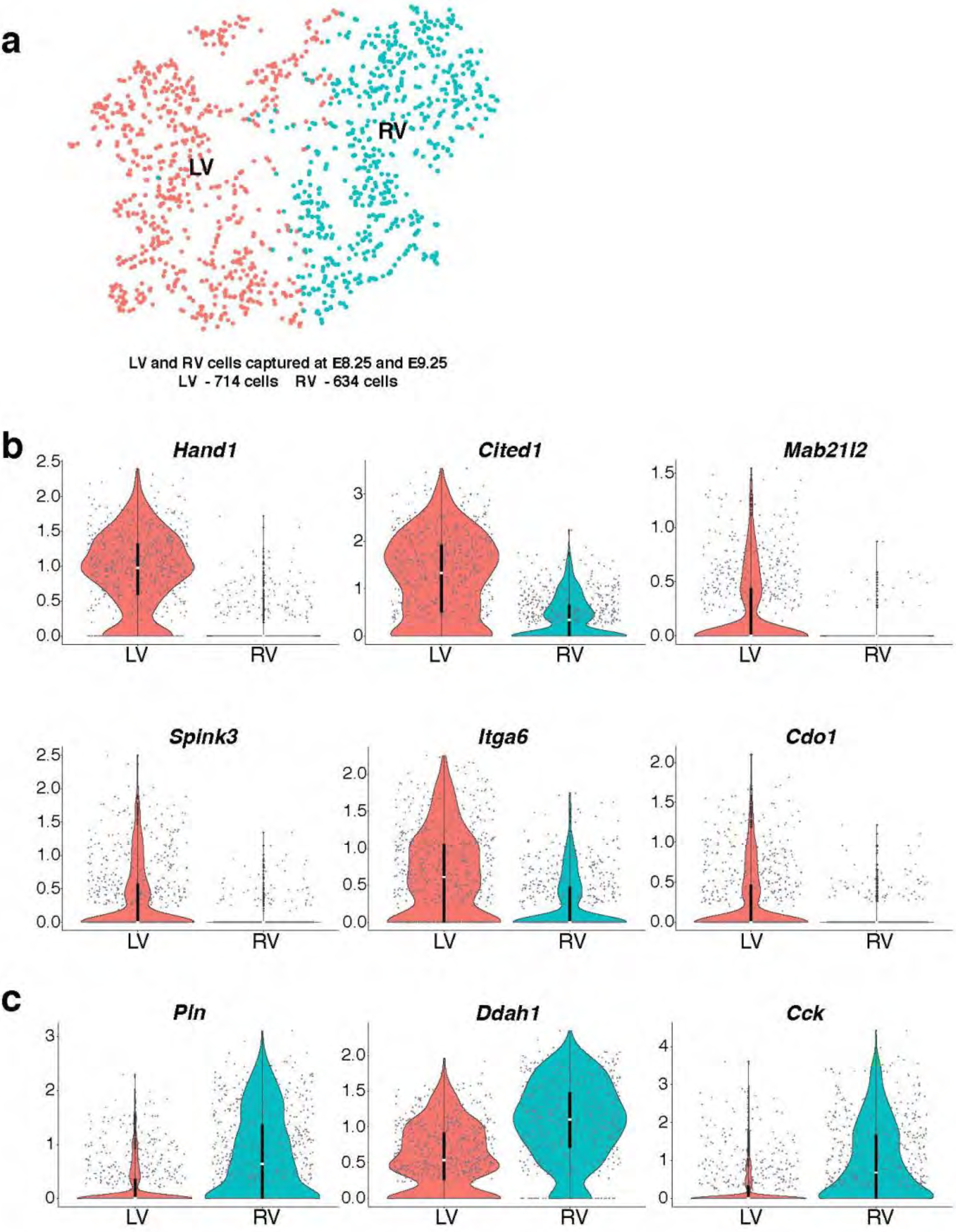
Analysis of LV and RV myocardium. **a**, tSNE plot of reclustered ventricle population from Fig 1b yields LV and RV myocardium subtypes. **b**, Violin plots of genes enriched in LV; all genes represented have an adjusted *p*-value < 1×10^−28^. **c**, Violin plots of genes enriched in RV; all genes represented have an adjusted *p*-value < 1×10^−52^. Summary statistics reported in violin plots: the centre white dot represents median gene expression and the central black rectangle spans the first quartile to the third quartile of the data distribution. The whiskers above or below the box indicate value at 1.5x interquartile range above the third quartile or below the first quartile.

**Extended Data Figure 3:**
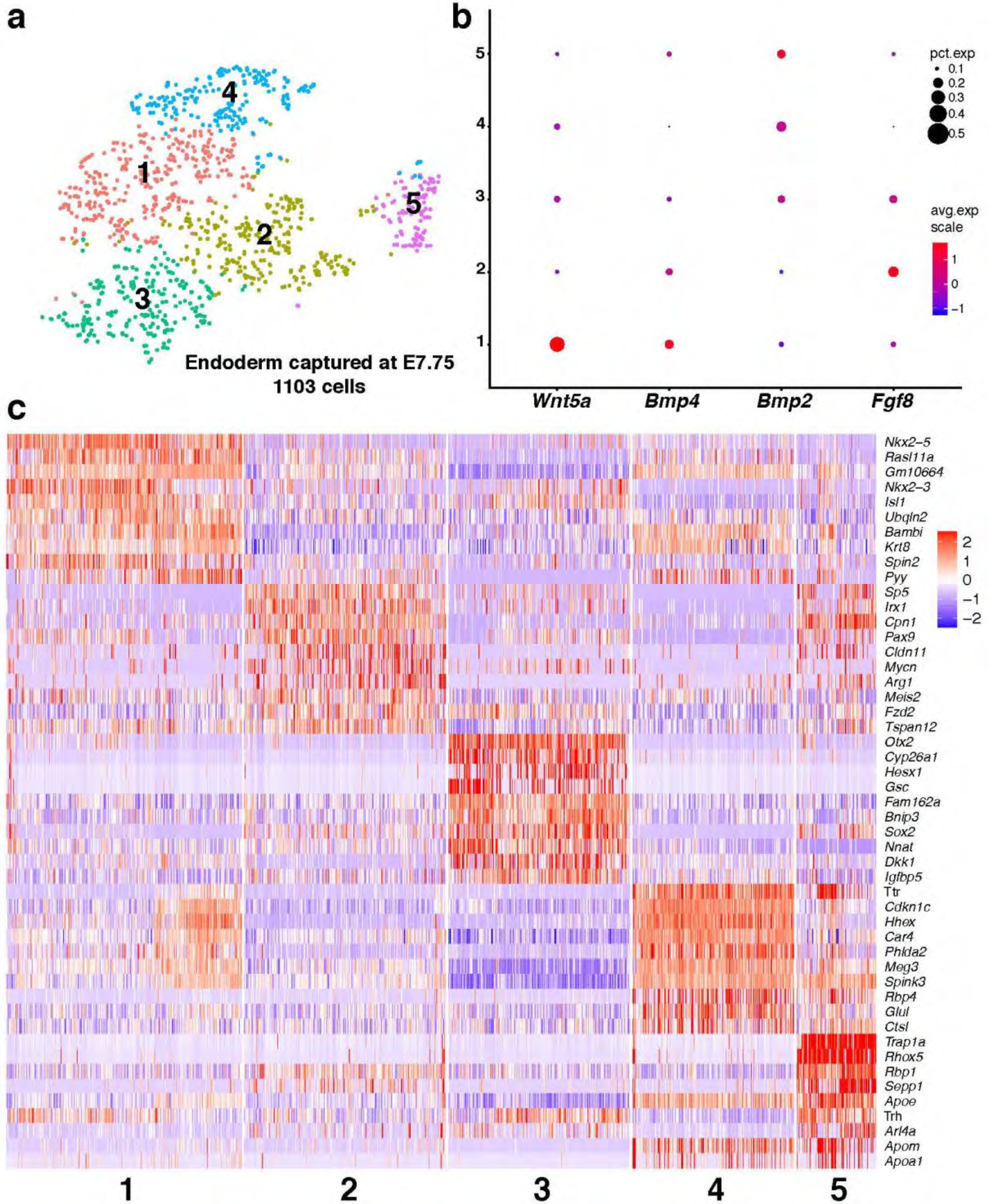
Identification of secreted factors from endoderm at E7.75. **a**, tSNE plot of endoderm populations captured at E7.75 colored by cluster. **b**, DotPlot highlighting expression patterns of known and novel endodermal secreted factors, *Fgf8, Bmp4, Bmp2*, and *Wnt5a*. **c**, Expression heatmap of the top ten marker genes of each endodermal population.

**Extended Data Figure 4:**
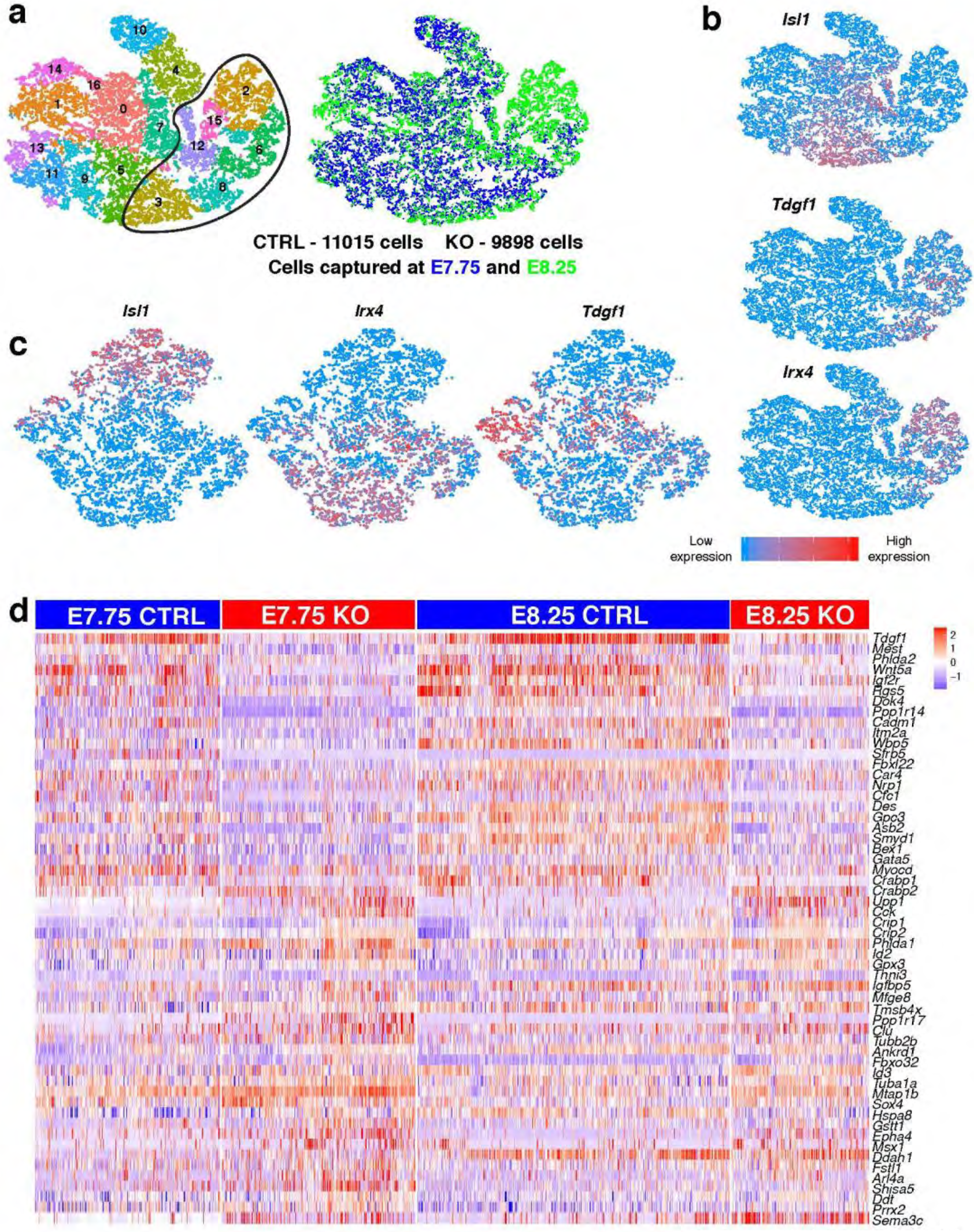
Transcriptional perturbation between CTRL and *Hand2-*null OFT/RV progenitors occurs as early as E7.75 and is exacerbated at E8.25. **a**, tSNE plot of all CTRL and *Hand2-*null progenitors captured at E7.75 and E8.25, colored by cluster and embryonic stage of collection. Circled region indicates reclustered populations represented in Fig. 3a. **b**, Expression pattern of *Isl1, Irx4* and *Tdgf1* that informed the selection of reanalyzed populations represented in Fig. 3a. **c**, Expression pattern of *Isl1, Irx4* and *Tdgf1* in tSNE plot represented in Fig. 3a. **d**, Expression heatmap of differentially expressed genes between CTRL and *Hand2-*null OFT/RV progenitors at E7.75 and E8.25. All genes have an adjusted *p*-value < 1×10^−4^.

**Extended Data Figure 5:**
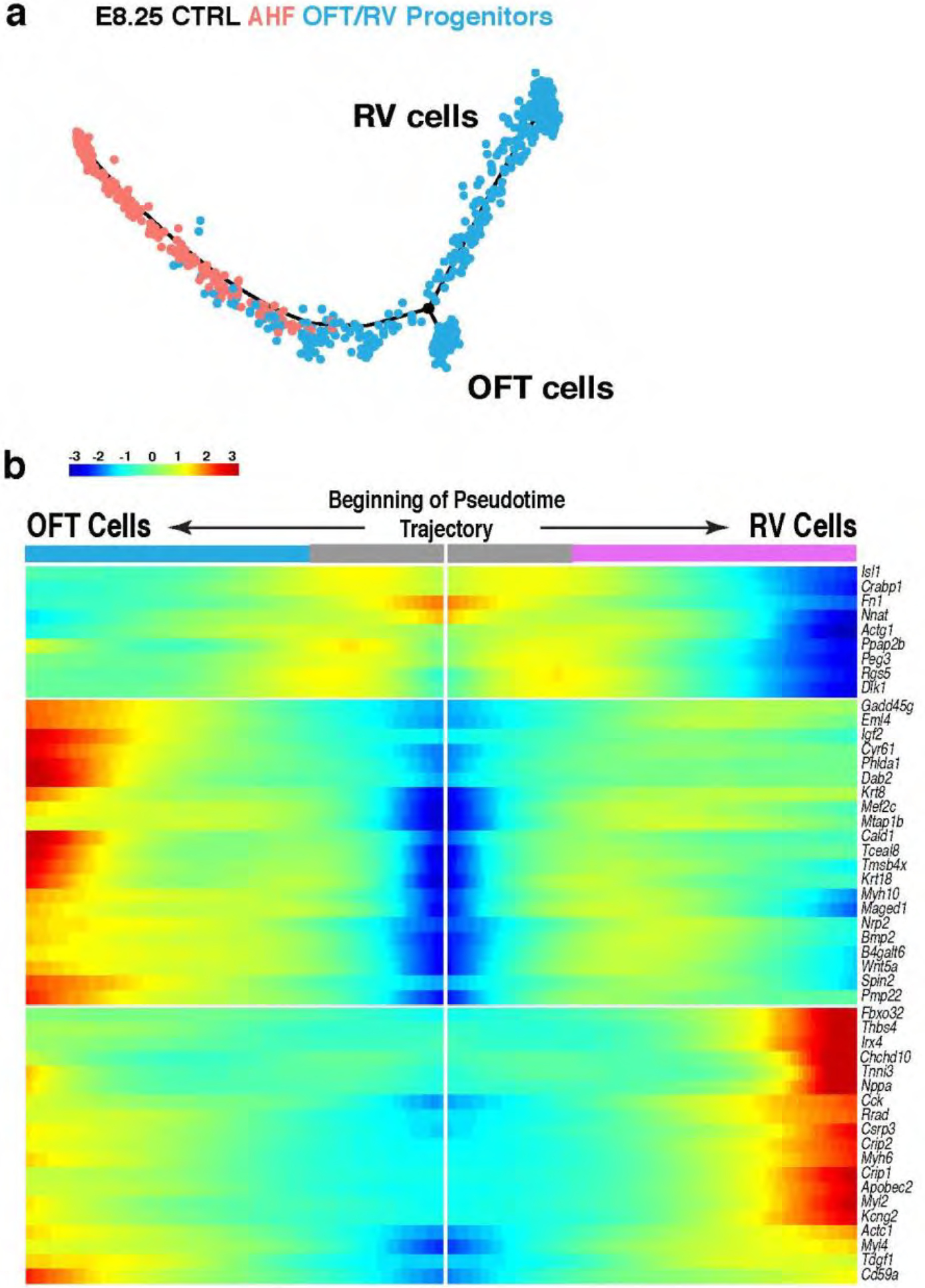
Pseudotime trajectory of CTRL and *Hand2*-null AHF and OFT/RV cells at E8.25. **a**, Pseudotime trajectory of only CTRL AHF and OFT/RV progenitors, colored by cluster. Black circle indicates branch point where trajectory bifurcates into two lineages. **b**, Branched heatmap of lineage-dependent gene expression identified by BEAM^32^ indicating the identities of each lineage as OFT and RV. Center of the heatmap indicates expression of genes at the start of the pseudotime trajectory. Reading the heatmap from the center to the right follows RV lineage through pseudotime; reading to the left follows the OFT lineage through pseudotime. Genes represented here have an adjusted *p*-value <10^−15^.

**Extended Data Figure 6:**
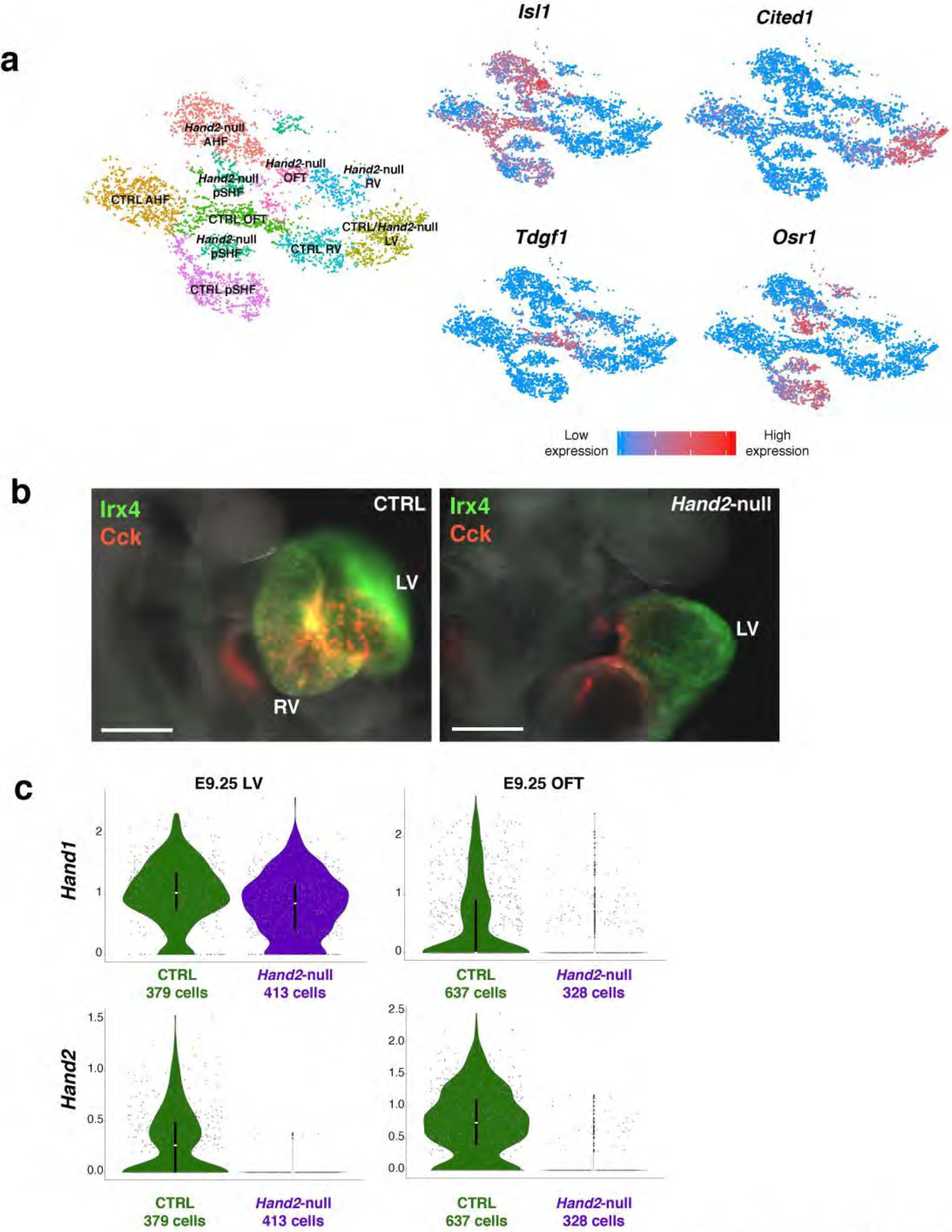
Identification of RV cells in *Hand2-*null embryos at E9.25 and expression dynamics of *Hand1* in *Hand2*-null embryos. **a**, Expression of *Isl1, Cited1, Tdgf1* and *Osr1* indicating presence of AHF, LV, OFT and pSHF populations, respectively, at E9.25. Red and blue indicate high and low expression, respectively. **b**, Expression of *Irx4* and *Cck* in CTRL and *Hand2*-null embryos at E9.0 indicating a migration defect in RV cells. Scale bar indicates 200 μm. **c**, Violin plots indicating expression dynamics of *Hand1* and *Hand2* in CTRL and *Hand2*-null LV and OFT cells. Adjusted *p*-value < 1×10^-^4. Summary statistics reported in violin plots: the centre white dot represents median gene expression and the central black rectangle spans the first quartile to the third quartile of the data distribution. The whiskers above or below the box indicate value at 1.5x interquartile range above the third quartile or below the first quartile.

